# Copper microRNAs govern the formation of giant feeding cells induced by the root knot nematode *Meloidogyne incognita* in *Arabidopsis thaliana*

**DOI:** 10.1101/2021.10.25.465754

**Authors:** Yara Noureddine, Martine da Rocha, Sébastien Thomine, Michaël Quentin, Pierre Abad, Bruno Favery, Stéphanie Jaubert-Possamai

**Affiliations:** INRAE, Université Côte d’Azur, CNRS, ISA, Sophia Antipolis, F-06903, France; Institute for Integrative Biology of the Cell (I2BC), UMR9198 CNRS/CEA/Univ. Paris Sud, Université Paris-Saclay, Gif-sur-Yvette, France

## Abstract

miR408 and miR398 are two conserved microRNAs which expression is activated by the SPL7 transcription factor in response to copper starvation. We identified these two microRNAs families as upregulated in *Arabidopsis thaliana* and *Solanum lycopersicum* roots infected by root-knot nematodes. These endoparasites induce the dedifferentiation of a few root cells and the reprogramming of their gene expression to generate giant feeding cells. By combining functional approaches, we deciphered the signaling cascade involving these microRNAs, their regulator and their targets. *MIR408* expression was located within nematode-induced feeding cells in which it co-localised with *SPL7* expression and was regulated by copper. Moreover, infection assays with *mir408* and *spl7* KO mutants or lines expressing targets rendered resistant to cleavage by miR398 demonstrated the essential role of the *SPL7/MIR408/MIR398* module in the formation of giant feeding cells. Our findings reveals how perturbation of plant copper homeostasis, *via* the *SPL7/MIR408/MIR398* module, governs the formation of nematode-induced feeding cells.

## Introduction

MicroRNAs are small non-coding RNAs that regulate the expression of protein-coding genes, mostly at the post-transcriptional level, in plants. They are major post-transcriptional regulators of gene expression in various biological processes, including plant development (Li and Zhang, 2016), responses to abiotic stresses (Barciszewska-Pacak et al., 2015), hormone signalling (Curaba et al., 2014), and responses to pathogens or symbiotic micro-organisms (Hoang et al., 2020; Weiberg and Jin, 2015). MicroRNAs and short interfering RNAs (siRNAs) were recently shown to play a key role in plant-pathogen crosstalk through trans-kingdom RNAi processes (Weiberg et al., 2013; Cai et al., 2018; Dunker et al., 2020). MicroRNAs are produced by the cleavage of long double-stranded RNA precursors by the DICER RNAse, generating 20-22 nucleotides miRNA duplexes composed of a mature (5P) and a complementary (3P) strand. One of the two strands is then incorporated into the RNA-induced silencing complex (RISC) to guide the major RISC protein, ARGONAUTE1 (AGO1), to the targeted messenger RNA (mRNA) on the basis of sequence complementarity. The hybridization of AGO1-bound miRNAs to their targets induces predominantly targeted mRNA degradation in plants, but it may also lead to an inhibition of mRNA translation (Axtell, 2013).

The miR408 and miR398 microRNA families are conserved so-called “copper microRNAs”, due to their involvement in the plant response to copper deficiency (Zhang et al., 2015; Yamasaki et al., 2007). Copper is an essential nutrient for plants, due to its function as a cofactor for many proteins. Copper proteins are involved in electron transport chains or function as enzymes in redox reactions. In plants, copper is involved in respiration, photosynthesis, ethylene perception, the metabolism of reactive oxygen species and cell wall remodelling (reviewed in Burkhead et al., 2009). Copper microRNAs accumulate in response to copper deficiency and their synthesis is repressed when copper concentrations are sufficiently high (Yamasaki et al., 2009). The underlying mechanism has been described in *Arabidopsis thaliana*, in which the regulation of *MIR408, MIR398B* and *MIR398C* by copper levels was shown to be mediated by the SQUAMOSA PROMOTER-BINDING PROTEIN-LIKE7 (SPL7) transcription factor (Yamasaki et al., 2009). The *A. thaliana* genome contains a single copy of *MIR408*, and three *MIR398* genes: *MIR398A, -B and -C*. At high copper concentrations, the DNA-binding activity of the SPL7 transcription factor is repressed, preventing the induction of transcription for downstream genes, such as *MIR408* or *MIR398B* and *MIR398C*, but not *MIR398A* (Sommer et al., 2011; Yamasaki et al., 2009). In the presence of low concentrations of copper, SPL7 activates the expression of copper-responsive microRNAs that target and repress the expression of genes encoding copper-binding proteins. These proteins are replaced by proteins that do not bind copper, to save copper resources for the functions for which this element is essential, such as photosynthesis (reviewed in Burkhead et al., 2009). For example, the mRNA for the cytosolic COPPER/ZINC (Cu-Zn) SUPEROXIDE DISMUTASE, CSD1, which can be replaced by an iron (Fe)-dependent SOD, is targeted by the copper microRNA miR398. In addition to their regulation as a function of copper levels through the activity of SPL7, *MIR408* and the *MIR398* family are also regulated by several environmental cues and abiotic stresses, such as light, which regulates *MIR408* activity through the ELONGATED HYPOCOTYL5 (HY5) or PHYTOCHROME INTERACTING FACTOR1 (PIF1) transcription factors in *A. thaliana* (Jiang et al., 2021), and salinity, oxidative and cold stresses, which have been shown to induce miR408 in *A. thaliana* (Ma et al., 2015), or cadmium treatment in *Brassica napus* (Fu et al., 2019). The miR408 and miR398 families have been widely analysed in plant responses to abiotic stresses, but little is known of their role in plant responses to biotic stresses. In sweet potato, *MIR408* has been associated with plant defences, as it is repressed by jasmonic acid (JA) and wounding, and miR408-overexpressing plants have attenuated resistance to insect feeding (Kuo et al., 2019). Moreover, miR398 has been shown to regulate cell death in response to the causal agent of barley powdery mildew, *Blumeria graminis* (Xu et al., 2014).

Root-knot nematodes (RKN) of the genus *Meloidogyne* are obligatory sedentary plant parasites capable of infesting more than 5,000 plant species (Abad and Williamson, 2010; Blok et al., 2008). The RKN larvae penetrate the roots, in which they induce the de-differentiation of five to seven parenchyma root cells and their reprogramming into the multinucleate, hypertrophied feeding cells that form the feeding site. These metabolically overactive feeding cells provide the nutrients required for RKN development (Favery et al., 2020). During the dedifferentiation of vascular cells and their conversion into ‘giant’ feeding cells, the cells surrounding the feeding site begin to divide again. The growth of the feeding cells and the division of the surrounding cells lead to a root swelling known as a gall. The feeding cell induction occurs in the first three days after root infection. Feeding cell formation can be split into two phases. Firstly, the cells undergo successive nuclear divisions coupled with cell expansion until ten days post infection (dpi) in *A. thaliana* (Caillaud *et al*., 2008). In the second phase, from 10 to 21 dpi, the successive nuclear divisions stop and the nuclei of the feeding cells undergo extensive endoreduplication (Wiggers et al., 1990; de Almeida Engler and Gheysen, 2013). The dedifferentiation of vascular cells and their conversion into giant cells result from an extensive reprogramming of gene expression in root cells, in response to RKN signals. In *A. thaliana*, the expression of approximately 10% of protein-coding genes is modified in galls induced by RKN (Cabrera et al., 2014; Yamaguchi et al., 2017; Jammes et al., 2005; reviewed in Escobar et al., 2011). The sequencing of small RNAs identified 24 mature microRNAs differentially expressed between *A. thaliana* galls induced by *M. incognita* and uninfected roots at 7 and 14 dpi (Medina et al., 2017). The miR408 and miR398 families of copper-responsive microRNAs were found to be upregulated in galls at 7 and/or 14 dpi.

In this article, we showed a conserved upregulation of these two microRNA families in *Arabidopsis* and tomato galls. Moreover, we found that the upregulation of miR408 in response to nematode was required for successful infection. Our findings highlighted a strong activity of *MIR408* promoter (*pMIR408*) in early galls that is i) driven by the modulation of environmental copper levels, ii) colocalised with strong *SPL7* expression. Moreover, we also demonstrated the involvement of this transcription factor in giant cell formation. In addition, we showed that the silencing of *CSD1* and *BLUE COPPER BINDING PROTEIN* (*BCBP*) transcripts by miR398 is involved in gall development. Finally, the watering of *Arabidopsis* with copper sulphate solutions at concentrations below the toxicity thresholds for nematode and plant development greatly decreased the RKN infection and impaired feeding cell development.

## Results

### The copper microRNA miR408 is crucial for the *Arabidopsis-Meloidogyne* interaction

Our previous analysis of microRNAs expressed in *Arabidopsis* galls induced by *M. incognita* revealed an upregulation of mature miR408 in *Arabidopsis* galls at 7 and 14 dpi, whereas miR398b/c was specifically upregulated in galls at 14 dpi. Sequencing of small RNAs from uninfected roots and galls of *Solanum lycopersicum* showed that these two microRNA families were also upregulated in tomato galls at 7 and 14 dpi (Table 1 and supplemental Table S1). Therefore, these microRNAs are among the very few conserved microRNAs which expression profile is conserved in *Arabidopsis* and tomato galls. We investigated the role of miR408 in gall development using two previously described *Arabidopsis* KO mutant lines: *miR408-1* and *miR408-2* (Maunoury and Vaucheret, 2011). The KO lines and corresponding wild-type plants were inoculated with *M. incognita* second stage juveniles (J2s) and their susceptibility was quantified by counting the galls and egg masses produced by adult females at the root surface. The two KO lines for *MIR408* had 40 to 50% fewer galls and egg masses than the wild type (*p*<0.05; Figure 1 and Supplemental Data Set S1). The roots of these KO lines were of similar weight and global architecture to those of wild-type plants (Supplemental Figure S1 and Supplemental Data Set S1). We then investigated the effect of the *MIR408* mutation on feeding site development, by comparing the area of feeding cells within galls collected from KO and wild-type plants (Figure 1A-C and Supplemental Data Set S1). Both KO mutants had a significantly smaller feeding site area than the wild type. Overall, these results demonstrate that *MIR408* is involved in feeding cell development in the *Arabidopsis*-nematode interaction, and that the lower susceptibility of the *miR408* KO lines is due to defects of feeding site formation.

**Figure 1:**
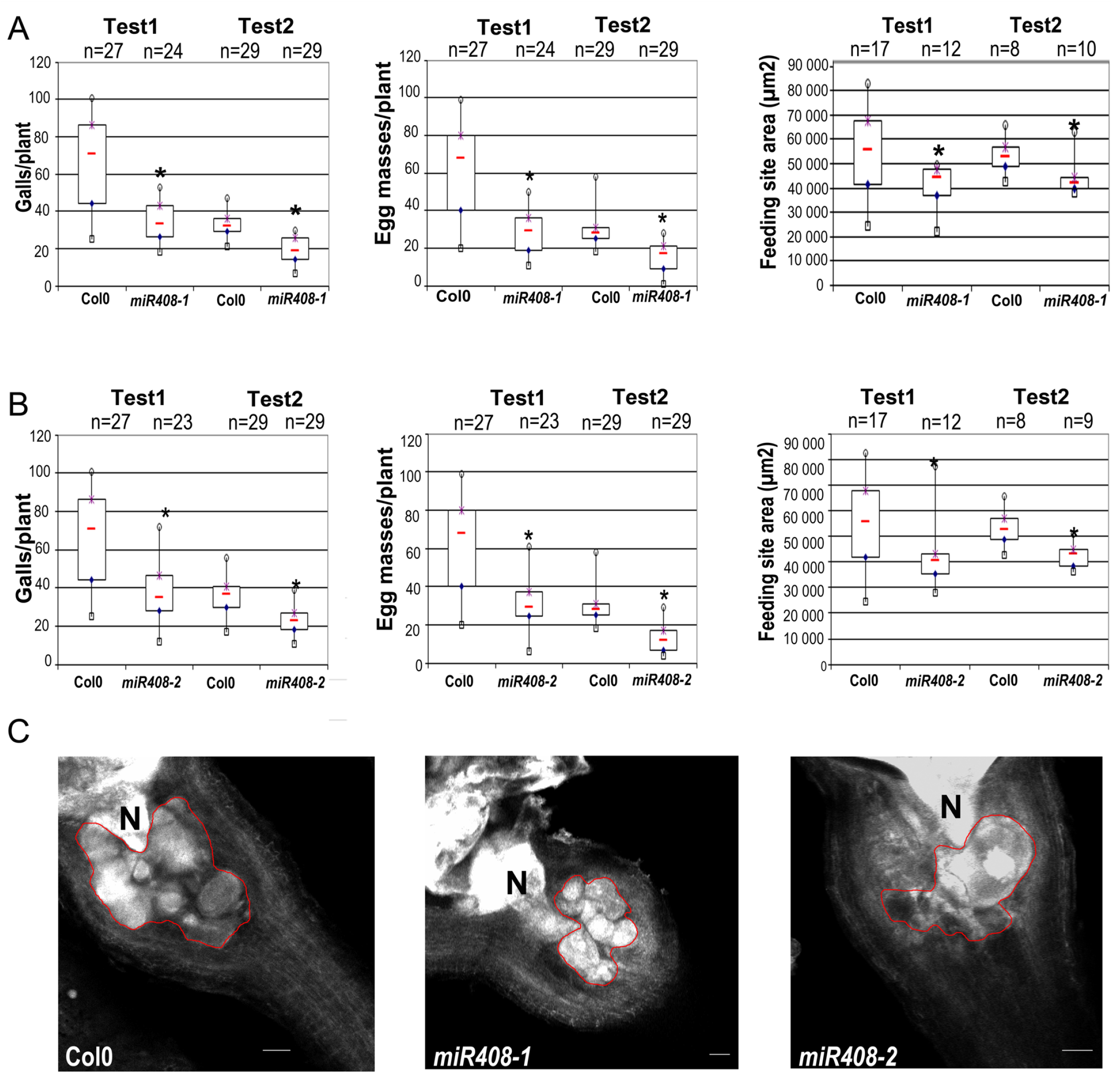
The miR408 KO lines were significantly less susceptible to M. incognita than the wild type. A-B, The susceptibility of the two miR408 KO lines, miR408-1 (A) & miR408-2 (B), and ColO wild type to M. incognita was evaluated by counting the number of galls and egg masses per plant in two independent infection assays in soil. The effect of miR408 mutation on the development of giant feeding cells was further evaluated by measuring the size of the feeding site produced in each KO line and comparing it to that in ColO. Galls were collected seven weeks post in vitro infection to measure the area (μm2) covered by the giant cells by the BABB clearing method (Cabrera et al., 2018). The impact of plant genotype was analysed in Mann and Whitney tests.*, P < 0.05. Open squares, minimum values; open circles, maximum values; red lines, median values; blue diamond, first quartile; purple star, third quartile. Bars 50μm.

**Table 1.**
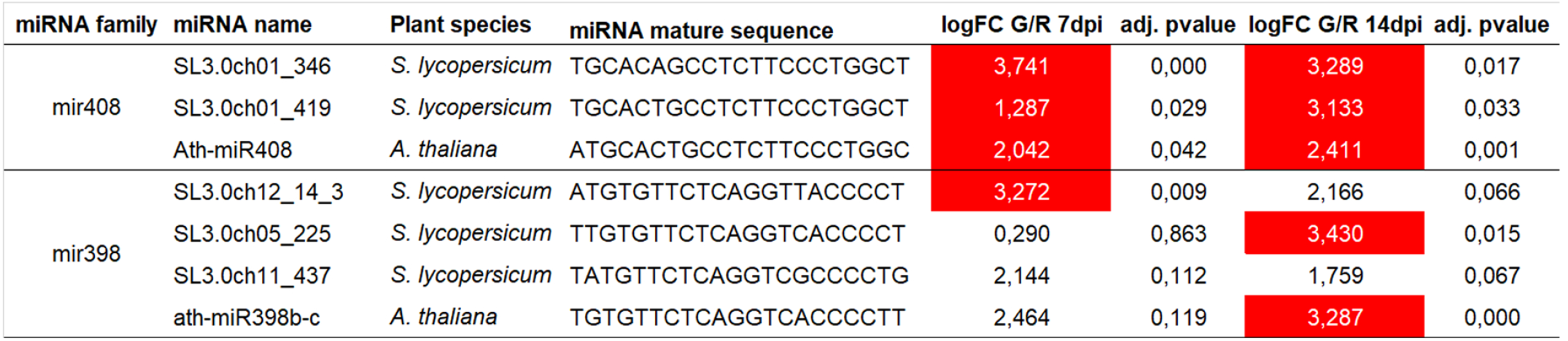
Expression profile of miRNAs of the miR408 and miR398 families in *Arabidopsis* and tomato galls.

We investigated the mechanisms by which miR408 regulates feeding cell formation in galls, by identifying the targets of miR408. The psRNA target algorithm (Dai et al., 2018) predicted 101 genes as putative targets of miR408 in *Arabidopsis* (Supplemental Data Set S2A). The expression profiles of these genes in galls at 7 dpi and 14 dpi were obtained from previous transcriptome analyses (Jammes *et al*. 2005). Only seven of the 101 putative targets were differentially expressed in galls at 7 and/or 14 dpi, and only two putative targets were repressed: a gene encoding a copper-binding protein, UCLACYANIN2 (*UCC2, At2g44790*), which is known to be cleaved by miR408 in senescing leaves and siliques (Thatcher et al., 2015), and a gene encoding a PHOSPHATASE 2G (*PP2CG1, At2g33700*) (Supplemental Data Set S2B).

### *MIR408* induction in galls is driven by modulation of copper level

We investigated the induction of *MIR408* in response to nematode infection, by inoculating plants expressing pMIR408::GUS with *M. incognita* J2s (Zhang and Li, 2013) *in vitro* in the presence of normal copper levels (0.1 μM CuSO_4_) or with a high copper concentration (Gamborg B5 plus 5 μM CuSO_4_). In the presence of normal concentrations of copper, we observed a strong GUS signal in developing galls at 3 and 7 dpi (Figure 2A-B). This signal had decreased in intensity by 14 dpi (Figure 2C) and disappeared completely from fully developed galls at 21 and 28 dpi (Supplemental Figure S2). On gall sections, the GUS signal was localised in the giant feeding cells and neighbouring cells, at the 3 dpi, 7 dpi and 14 dpi time points (Figure 2D-F). By contrast, in plants grown in the presence of high copper concentrations, the GUS signal was much weaker in galls at 3 and 7 dpi (Figure 2G-H), and undetectable in galls at 14 dpi (Figure 2I). The repression of *MIR408* expression in galls by high copper concentrations indicates that *MIR408* expression within *Arabidopsis* galls is regulated by modulation of copper levels.

**Figure 2.**
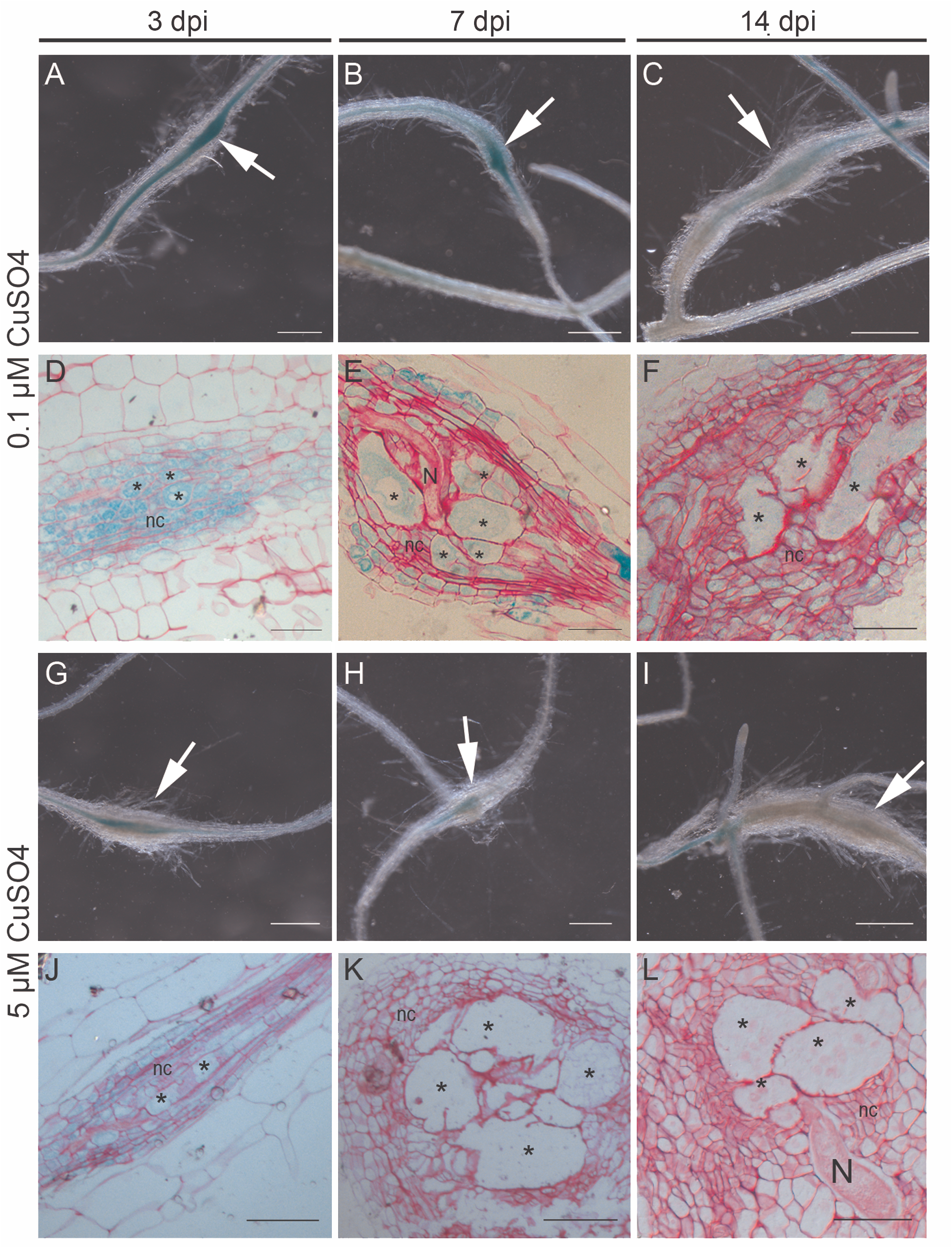
Copper modulates MIR408 promoter activity in galls. A-H, The activity of the MIR408 promoter was analysed in galls induced by M. incognita in Arabidopsis expressing the pMIR4O8::GUS construct grown in the presence of normal concentrations of copper (0.1 μM CuSO_4_)(A-F) or in the presence of high concentrations of copper (5.0 μM CuSO_4_)(G-L). A-C, a strong GUS signal was observed in galls 3 days post infection (dpi) (A), 7 dpi (B) and 14 dpi (C) in plants grown with 0.1 μM CuSO_4_. D-F, Section of gall at 3 dpi (D), 7dpi (E) and 14 dpi (F) showing the GUS signal in giant cells and in the cells surrounding the giant cells. G-l, a weaker GUS signal was observed in galls from plants grown with 5.0 μM CuSO_4_ analysed at 3 dpi (G) and 7 dpi (H) and no GUS signal was observed in galls at 14 dpi (I). J-L, Section of gall at 3 dpi (J), 7dpi (K) and 14 dpi (L). Galls are indicated with an arrow; N, nematode; (*) giant feeding cells; nc, neighbouring cells. Bars 500 μm (A-C; G-l) or 50 μm (D-F; J-L).

### SPL7 is an activator of *MIR408* transcription in galls

The regulation of *MIR408* by modulation of copper levels has been shown to be mediated by the SPL7 transcription factor (Zhang et al., 2014; Bernal et al., 2012). Transcriptome analyses from *Arabidopsis* galls showed that *SPL7* was expressed in galls at 7, 14 and 21 dpi (Jammes et al., 2005). We further investigated the expression of *SPL7* within galls, by inoculating a *pSPL7::GUS Arabidopsis* line (Araki et al., 2018) with *M. incognita* in the presence of a normal copper concentration. *SPL7* promoter activity was observed within the gall from 3 to 14 dpi, with lower levels at 14 dpi (Figure 3A-C). As observed for *pMIR408::GUS*, sections of*pSPL7::GUS* galls revealed a GUS signal in the giant feeding cells and neighbouring cells (Figure 3D). We investigated the putative function of *SPL7* in plant responses to RKN, by inoculating the *Arabidopsis spl7* KO mutant described by Zhang *et al*. (2014) with *M. incognita* J2s. *SPL7* knockout led to the production of smaller numbers of galls and egg masses per plant than were observed for the wild-type (Figure 3E-F and Supplemental Data Set S3). This knockout had no effect on root weight (Supplemental Figure S3 and Supplemental Data Set S3). Measurements of the area of the feeding site within galls revealed defects of feeding site formation in the *spl7* KO mutants, resulting in smaller giant cells than were observed in wild-type plants (Figure 3G and Supplemental Data Set S3). Overall, these results demonstrate the requirement of miR408 and *SPL7* for the development of giant cells. The upregulation of mature miR408 observed in galls suggest an induction of *MIR408* expression driven by SPL7 due to a decrease in copper availability within the gall.

**Figure 3.**
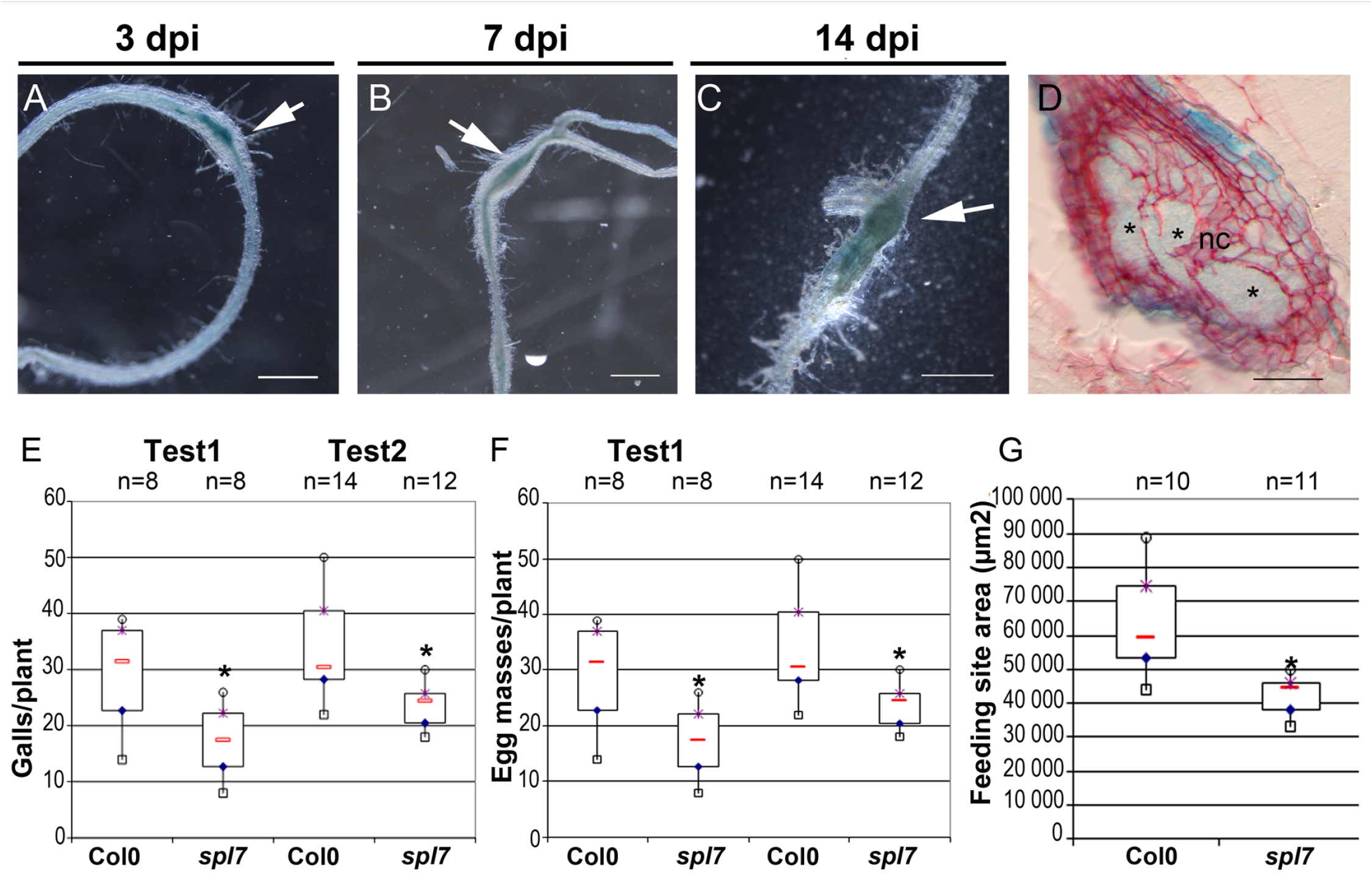
SPL7 is induced and required for M. incognita infection and giant cell formation in Arabidopsis. A-D, The activity of the SPL7 promoter (pSPL7) was studied in galls induced by M. incognita from A. thaliana expressing the pSPL7::GUS construct, at 3 days post inoculation (dpi)(A), 7 dpi (B) and 14 dpi (C). D, GUS activity was observed within 5.0 μm-thick gall sections at 7 dpi. E-G, the KO spl7 line (SALKO93849c) was infected with M. incognita J2. This line was significantly less susceptible to RKN than ColO, as shown by the smaller mean number of galls (E) and egg masses (F) per plant in two infection assays. (G), the effect of spl7 mutation on the development of feeding cells was further evaluated by measuring the size of the feeding site produced. Galls were collected seven weeks post in vitro infection for measurement of the area (μm2) covered by the giant cells, by the BABB clearing method (Cabrera et al., 2018). Mann-Whitney tests were performed for statistical analysis in each experiment; significant differences relative to Col-0: *, P < 0.05; Open squares, minimum values; open circles, maximum values; red lines, median values; blue diamond, first quartile; purple star, third quartile. Galls are indicated with an arrow; (*) giant feeding cells; nc, neighbouring cells. Bars 50 μm (A-C) or 500 μm (D).

### micro398, a second copper-responsive microRNA family involved in the *Arabidopsis-Meloidogyne* interaction

MiR408 is not the only copper-responsive microRNA differentially expressed in galls. The expression of *MIR398B* and *MIR398C*, from the conserved miR398 family, has also been shown to be induced in response to copper deficiency, via SPL7 activity (Araki et al., 2018). We previously described an induction of the mature miR398b and miR398c in *Arabidopsis* galls at 14 dpi (Medina et al., 2017). Three targets of the miR398 family have been biologically validated: the *At1g08830* and *At2g28190* transcripts encoding two copper superoxide dismutases, *CSD1* and *CSD2*, respectively, and the *At5g20230* transcript encoding the blue copper binding protein (BCBP), identified as a non-canonical target of miR398 (Brousse et al., 2014). We investigated the role of miR398 further, by infecting *Arabidopsis* lines expressing modified versions of *CSD1* (*mcsd1*) or *BCBP* (*mbcbp*) mRNAs rendered resistant to cleavage by miR398 (Brousse et al., 2014; Beauclair et al., 2010). The target mRNA levels is therefore artificially increased in these plants. The prevention of *CSD1* transcript cleavage by miR398 had no effect on root weight (Figure S4), but led to lower levels of nematode infection, with the mutant having less galls and egg masses than the wild type (Figure 4 and Supplemental Data Set S4). The *mbcbp* line also had fewer egg masses than the wild-type (Figure 4 and Supplemental Data Set S4). No defect of feeding cell formation, such as slower feeding cell growth, was observed in either of these lines (Figure 4 and Supplemental Data Set S4). The specific decrease in egg mass production by females provides evidence for a role for miR398 in the functionality of feeding cells, although the mutations in the *mcsd1* and *mbcbp* lines did not affect feeding site size. These findings demonstrate that the cleavage of *CSD1* and *BCBP* transcripts by miR398 is required for plant-RKN interaction.

**Figure 4.**
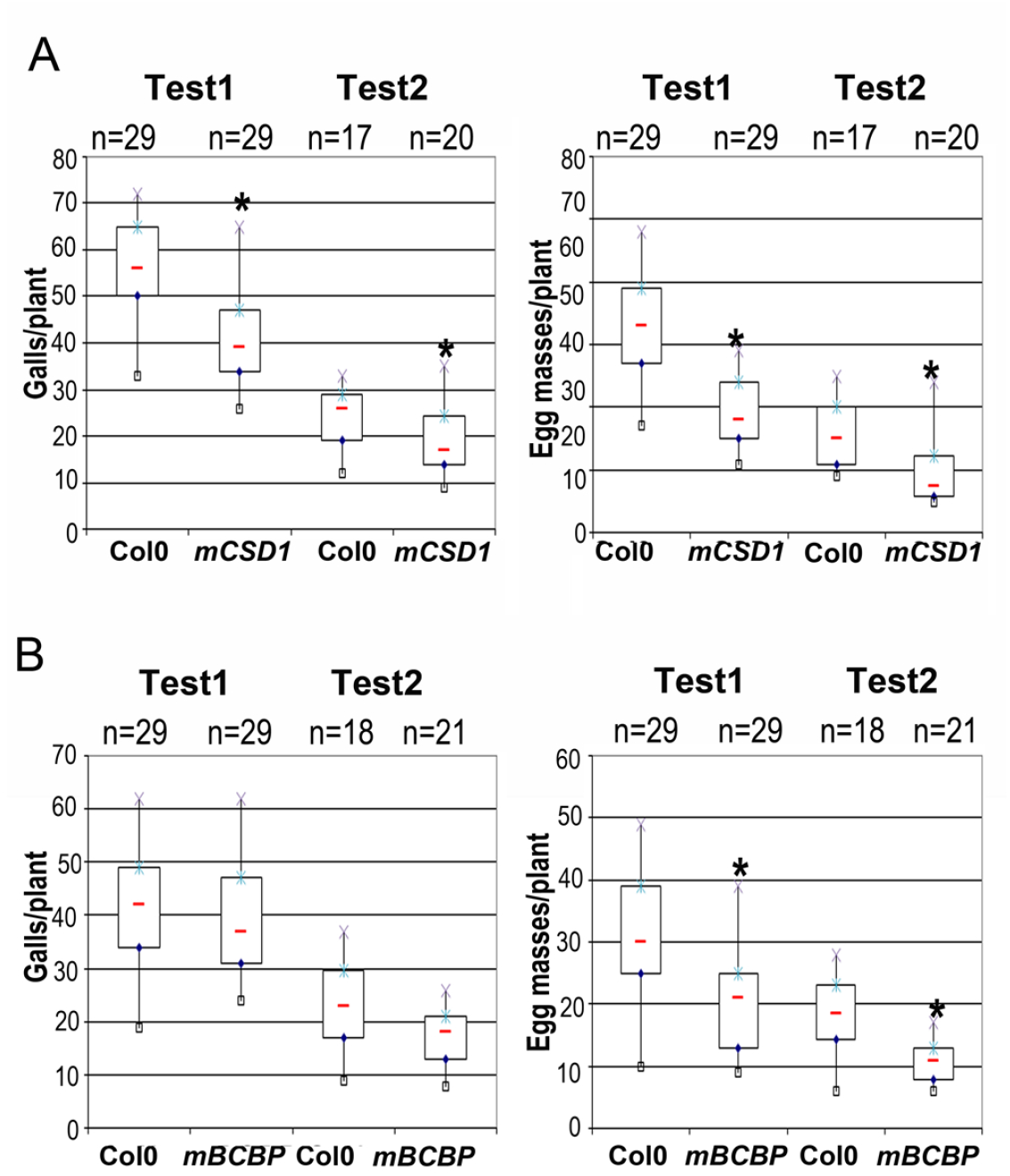
The miR398-resistant mcsdl and mbcbp mutant lines had smaller numbers of egg masses. The susceptibility of the mcsdl and mbcbp lines and of wild-type ColO plants was evaluated by counting the number of galls and egg masses per plant in two independent infection assays in soil. The impact of the plant genotype on the number of galls and egg masses relative to ColO was analysed in Mann and Whitney statistical tests.*, P < 0.05. Open squares, minimum values; open circles, maximum values; red lines, median values; blue diamond, first quartile; purple star, third quartile.

### Modulation of copper levels is essential for plant-RKN interaction

To study further the effect of copper on nematode infection, we analysed the direct effects of copper on nematode survival and gall development. Free-living *M. incognita* J2s were incubated in several concentrations of copper sulphate (50 μM to 2 mM) used in previous studies assessing the effect of copper on plant development (Schulten et al., 2019). As a negative control, J2s were incubated in tap water. Living J2 counts after 24 hours in the copper sulphate solution showed that copper was non-toxic at a concentration of 50 μM (Supplemental Figure S5 and Supplemental Data Set S5). By contrast, toxic effects were observed for all other concentrations tested (0.5 mM, 1 mM and 2 mM). We then analysed the effect of copper on gall formation in Col0 and *pMIR408::GUS* plants grown in soil watered with 50 μM CuSO_4_. We also minimised J2 exposure to copper in the soil, by beginning to water plants 50 μM CuSO_4_ two days after inoculation, after the J2s had already penetrated the roots. Watering with 50 μM CuSO_4_ repressed pMIR408 activity, confirming the effects of such treatment in galls (Supplemental Figure S6 and Supplemental Data Set S6). Watering with 50 μM CuSO_4_ had no visible effect on root weight and architecture (Supplemental Figure S7 and Supplemental Data Set S7), but it resulted in a strong and significant decrease in the number of galls and egg masses relative to control plants watered with tap water (Figure 5). These results demonstrate that perturbation of plant copper homeostasis governs the formation plant-RKN interaction.

**Figure 5.**
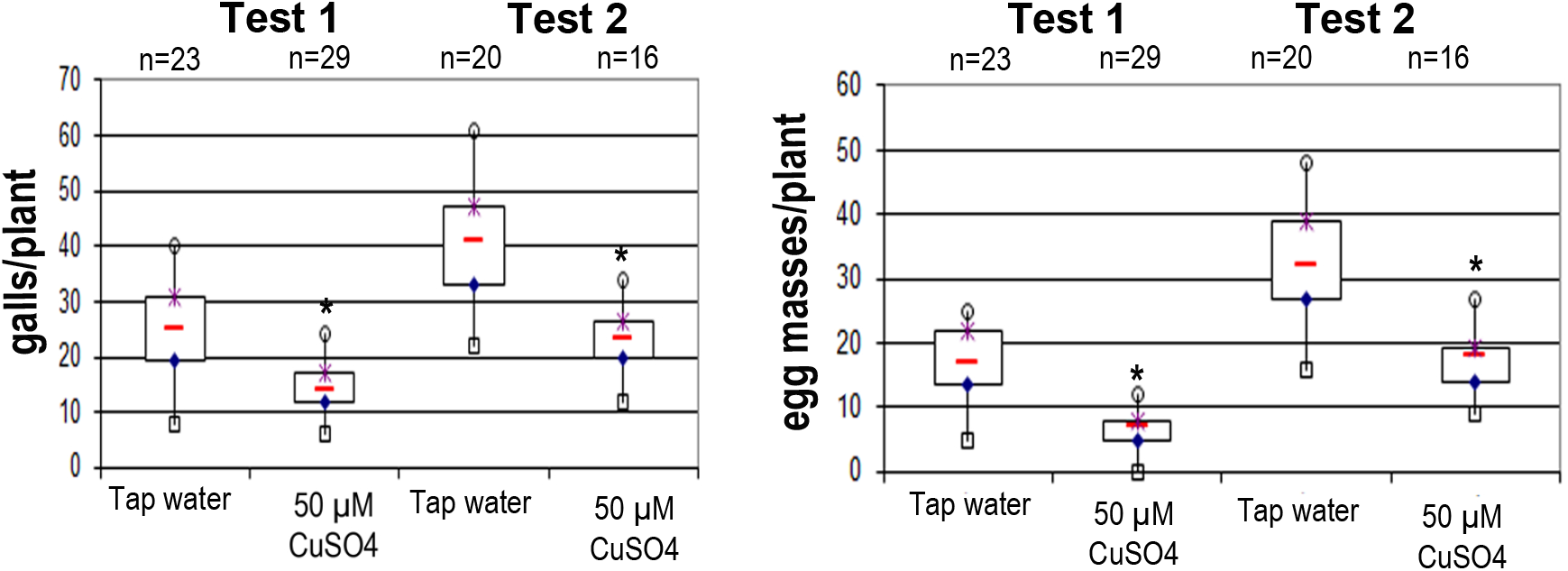
Plants watered with copper sulphate solution were significantly less susceptible to root knot nematodes. The effect of physiological concentrations of copper on M. incognita infection was evaluated by counting galls and egg masses in ColO plants watered with a copper sulphate solution at a non-toxic concentration (50 μM) and comparing the results to those for ColO watered with tap water. The impact of plant genotype on the numbers of galls and egg masses relative to ColO was analysed in Mann and Whitney statistical tests.*, P < 0.05. Open squares, minimum values; open circles, maximum <alues; red lines, median values; blue diamond, first quartile; purple star, third quartile.

## Discussion

RKN induce the formation of similar giant feeding cells in thousands of plant species. The conservation of the ontogeny and phenotype of nematode-induced feeding cells between species, strongly suggests that the plant molecular mechanisms manipulated by RKN are widely conserved across the plant kingdom. Previous transcriptome analyses on various plant species have shown that the development of galls in roots infected by RKN is associated with a massive reprogramming of gene expression (reviewed in Escobar et al., 2011). MicroRNAs are small non-coding RNAs that regulate gene expression at the post-transcriptional level, and some microRNA families, such as the miR156 and miR167 families, are widely conserved in plants (Chavez Montes Nature Communications 2014). The role for microRNAs in controlling gene expression during the formation of galls was recently reported in *Arabidopsis* (reviewed in Jaubert-Possamai et al., 2019), for the conserved microRNAs miR390, miR172 and miR159 (Escobar et al., 2015; Medina et al., 2017).

### miR408 and miR398: two copper-responsive microRNA families activated in *Arabidopsis* galls induced by *M. incognita*

A previous analysis of the levels of mature microRNAs in galls at 7 and or 14 dpi and in uninfected roots showed that mature mi408 and mir398b/c were induced in galls in response to *M. incognita* (Medina et al., 2017). A combination of *in silico* predictions of the transcripts targeted by miR408 and previous transcriptional analyses of galls and uninfected roots identified two putative targets downregulated in galls the genes the *UCLACYANIN-2* (*UCC2*) and the *PHOSPHATASE* (*PP2CG*). The cleavage of *UCC2* transcripts by miR408 has been biologically validated in *Arabidopsis* and rice (Thatcher et al., 2015; Zhang et al., 2017). Moreover, several targets of miR398 have been biologically validated, including the cytosolic *CSD1* and chloroplastic *CSD2*, and the non-canonical target *BCBP* (Beauclair et al., 2010; Brousse et al., 2014). Our analysis, thus, identified several biologically validated and conserved targets that may be considered robust candidates for mediating the functions of miR398b/c and miR408 in galls.

The inactivation of miR408 in T-DNA mutant lines, or of miR398b/c function in transgenic plants expressing mutated *CSD1* or *BCBP* resistant to miR398 cleavage, led to decreases in both the parameters used to assess parasitic success (the number of galls and the number of egg masses per root). The smaller number of galls in the mutant lines demonstrates the involvement of the miR398 family and miR408 in the early plant response to RKN. Moreover, the smaller feeding sites observed in the two *miR408* KO lines and the *spl7* mutant demonstrate that this miR408 and SPL7 are involved in the formation of the giant cells, which are essential for nematode growth and development. Females are unable to develop normally if the feeding cells are too small, as already reported in some *Arabidopsis* mutants, such as lines with a knockout of *PHYTOSULFOKINE RECEPTOR1* (*PSKR1*) (Rodiuc et al., 2016). Only a few genes and plant functions have been demonstrated to be essential for the formation of giant feeding cells (Favery *et al*., 2020). In the absence of changes in giant cell size in the *mcsd1* and *mbcbp* mutants, we hypothesise that the miR398-regulated *CSD1* and *BCBP* genes may play a role in giant cell functioning, potentially in reactive oxygen species (ROS)-related redox regulation and signalling (Zhao et al., 2020). Further studies will be required to determine their precise roles in the plant-RKN interaction.

### SPL7 is a regulator of miR408 and miR398 in galls

In *A. thaliana*, it has been shown that *MIR408, MIR398B* and *MIR398C* are activated by the same transcription factor, SPL7, the activity of which is dependent on copper levels (Yamasaki et al., 2009; Araki et al., 2018). We confirmed the activity of the *SPL7* promoter and *MIR408* in feeding cells and neighbouring cells. The co-expression of *SPL7* and *MIR408* within developing galls, and the similar nematode infection phenotype, with feeding site formation defects, strongly suggest that *SPL7* is responsible for activating *MIR408* transcription in galls, as already reported in leaves and the root vasculature (Yamasaki et al., 2009; Araki et al., 2018). We therefore hypothesised that the expression of *MIR408* and *MIR398B* and -*C* is activated by the SPL7 transcription factor in response to a decrease in copper concentration within galls. Other transcription factors, such as HY5 and PIF1, which are known to regulate the expression of *MIR408* in response to light stress (Zhang et al., 2014; Jiang et al., 2021), are expressed in galls and could also play a role in the regulation of *MIR408* expression. However, the strong repression of *MIR408* by excess copper observed in galls suggests that *MIR408* upregulation in galls is predominantly driven by copper and SPL7.

### Modulation of copper levels, a key conserved factor for gall formation

Infection assays with *A. thaliana* plants watered with a copper sulphate solution at a concentration non-toxic for plants and nematodes, showed a strong decreased RKN infection rates and resulted in defective feeding site formation. Together with the upregulation in galls of two microRNA families known to be induced by copper deficiency, the miR408 and miR398 families, and the regulation of *MIR408* expression by copper, this finding suggests that copper content decreases in the galls induced by RKN infection. This hypothesis is supported by the downregulation of the *COPT2* gene, encoding a copper importer, in the *Arabidopsis* gall transcriptome (Jammes et al., 2005). Assays in RKN-infected susceptible tomato roots have also demonstrated a decrease in copper concentration (Lobna et al., 2017).

The *SPL7*/*MIR408-UCC2/MIR398-CSD1* copper signalling cascade may be a key factor in gall formation, conserved across the plant kingdom. MiR408 and miR398 have been identified in more than 40 plant species (Griffiths-Jones et al., 2007) and *SPL7* is widely conserved throughout the plant kingdom (Yamasaki et al., 2009). Moreover, the targeting of *UCLACYANIN* by miR408 and of *CSD1* by miR398 is conserved in both dicotyledonous and monocotyledonous plants (Zhang et al., 2017; Thatcher et al., 2015). A role for UCLACYANINS in the formation of a lignified nanodomain within the Casparian strips known to form an endodermal barrier in *Arabidopsis* roots has recently been described (Reyt et al., 2020). Casparian strip defects have been observed in the endodermis bordering the giant cell area within sorghum galls induced by *M. naasi* (Ediz and Dickerson, 1976). Moreover, *Arabidopsis* mutants with disrupted Casparian strips are particularly susceptible to RKN (Holbein et al., 2019). The infection of plants with nematodes may, therefore, provides a unique model for investigating the role of copper modulation, *via* miR408 and its *UCLACYANIN2* target, in the formation of Casparian strips.

## Methods

### Biological material, growth conditions and nematode inoculation

Seeds of *A. thaliana* Col0 and mutants *miR408-1*(SALK_038860), *miR408-2* (SALK_121013.28.25.n), *spl7* (SALK_093849), *mcsd1* and *mbcbp* (Beauclair et al., 2010), *pmiR408::GUS* (Zhang et al., 2013) and *pSPL7::GUS* (Yamasaki et al., 2009) were surface-sterilised and sown on Gamborg B5 medium agar plates (0.5 x Gamborg, 1% sucrose, 0.8% agar, pH 6.4). The plates were incubated at 4°C for two days, then transferred to a growth chamber (20°C with an 8 h light/ 16 h darkness cycle). *M. incognita* strain “Morelos” was multiplied on tomato plants in a growth chamber (25°C, 16 h light/8 h darkness). For the RKN infection of plants in soil, two-week-old plantlets grown *in vitro* were transferred to a mixture of 50% sand (Biot B5)/50% soil in a growth chamber (21°C, 8 h light/16 h darkness). For studies of the effect of copper on gall development on plants *in vitro*, *Arabidopsis* plantlets were sown and cultured *in vitro*, as described above, on Gamborg B5 medium supplemented with 50 μM CuSO_4_.

### Root knot nematode infection assay

For nematode infections *in vitro*, J2s were surface-sterilised with HgCl_2_ (0.01 %) and streptomycin (0.7 %), as described by Caillaud and Favery (2016). We inoculated each 25-day-old seedlings grown individually *in vitro* with 200 sterilised J2s resuspended in Phytagel (5 %). Infection assays were performed on *Arabidopsis* mutants and a wild-type ecotype in soil. We inoculated 20 to 30 two-month-old plantlets with 150 J2s per plant and incubated them in a growth chamber (21°C, 8 h light/16 h darkness). Seven weeks after infection, the roots were collected, washed in tap water and stained with eosin (0.5 %). Stained roots were weighed and galls and egg masses were counted on each root under a binocular microscope. Mann and Whitney tests (2.5 %) were performed to determine the significance of the observed differences in the numbers of egg masses and galls per root.

### Small RNA sequencing from galls and uninfected tomato roots

Biological material, RNA extraction, small RNA sequencing, read mapping and statistical analysis are presented as supplemental material.

### BABB clearing

Feeding site development was evaluated by the BABB clearing method described by Cabrera et al., (2018). Briefly, the area occupied by the giant cells was measured on galls collected 14 dpi, cleared in benzyl alcohol/benzyl benzoate (BABB) and examined under an inverted confocal microscope (model LSM 880; Zeiss). Zeiss ZEN software was used to measure the area occupied by the giant cells in each gall, on two biological replicates. Data were analysed in Mann and Whitney tests.

### Copper treatment

*M. incognita* eggs were collected as previously described (Caillaud and Favery, 2016) and placed on a 10 μm-mesh sieve for hatching in tap water. Free-living J2s were collected from the water with a 0.5 μm-mesh sieve. We evaluated the toxicity of copper to J2s by incubating freshly hatched J2s in solutions of copper sulphate of various concentrations for 24 hours. The numbers of living or dead J2s were then determined by counting under a binocular microscope. We investigated the effects of copper on plant-nematode interactions in *Arabidopsis* Col0 grown in soil. *Arabidopsis* Col0 plantlets were prepared and inoculated as previously described for in-soil infection. Half the plants were watered with 50 μM CuSO_4_ two days after inoculation with J2s and then once per week for the next seven weeks. Control Col0 plants were watered with tap water in place of copper sulphate solution, at the same frequency. Seven weeks after inoculation, the plants were collected, their roots were washed and weighed, and the numbers of galls and egg masses on the roots were counted, as described above.

### Studies of promoter-GUS fusion gene expression

We localised the promoter activity of *MIR408* and *SPL7* in *A. thaliana* lines expressing various fusions of the GUS reporter gene to promoters from these genes (ref & Supplemental Table S7). We inoculated 21-day-old seedlings in soil and *in vitro*, as described above. We collected inoculated roots and washed them in water, 3, 7, 14 and 21 dpi. GUS staining was performed as previously described (Favery *et al*., 1998), and the roots were observed under a Zeiss Axioplan 2 microscope. Stained galls were dissected, fixed by incubation in 1% glutaraldehyde and 4% formaldehyde in 50 mM sodium phosphate buffer pH 7.2, dehydrated, and embedded in Technovit 7100 (Heraeus Kulzer, Wehrheim, Germany), according to the manufacturer’s instructions. Sections were cut and mounted in DPX (VWR International Ltd, Poole, UK), and observed under a Zeiss Axioplan 2 microscope (Zeiss, Jena, Germany).

### Bioinformatic analysis

We used psRNA target with default parameters for the prediction of miR408 targets (Dai, Zhuang and Zhao 2018).

## Author contributions

Y.N., B.F and S.J.P. designed the study and performed the experimental work. All authors analysed and discussed the data. S.J.P and B.F. designed the study and wrote the manuscript. M.Q. and P.A. participated in the writing of the manuscript. S.T. participated in the copper studies. MdR analysed NGS data. The author responsible for distributing materials integral to the findings presented in this article, in accordance with the policy of the journal (https://academic.oup.com/plcell) is: Stephanie Jaubert Possamai.

## Acknowledgements

We would like to thank Dr. Nicolas Bouché (INRAE Versailles, France) for helpful discussions and for providing the *mbcbp* and *mcsd1 Arabidopsis* lines. We would like to thank Dr. Lei Li (Beijing University, China) for providing the *pMIR408:GUS Arabidopsis* line. We would also like to thank Dr. Toshiharu Shikanai (Kyoto University, Japan) for providing the *pSPL7:GUS* and ko *spl7 Arabidopsis* line. The microscopy work was performed at the SPIBOC imaging facility of Institut Sophia Agrobiotech. We thank Dr Olivier Pierre and the entire team of the platform for assistance with microscopy. This work was funded by the INRA SPE department and the French Government (National Research Agency, ANR) through the ‘Investments for the Future’ LabEx SIGNALIFE: programme reference #ANR-11-LABX-0028-01 and IDEX UCAJedi ANR-15-IDEX-0, and by the French-Japanese bilateral collaboration programme PHC SAKURA 2019 #43006VJ. Y.N. was supported by a doctoral fellowship from Lebanon (Municipal Council of Aazzée).

